# Cortical thickness estimation in individuals with cerebral small vessel disease, focal atrophy, and chronic stroke lesions

**DOI:** 10.1101/2020.08.04.236760

**Authors:** Miracle Ozzoude, Joel Ramirez, Pradeep Raamana, Melissa F. Holmes, Kirstin Walker, Christopher J.M. Scott, Maged Goubran, Donna Kwan, Maria C. Tartaglia, Derek Beaton, Gustavo Saposnik, Ayman Hassan, Jane Lawrence-Dewar, Dariush Dowlatshahi, Stephen C. Strother, Sean Symons, Robert Bartha, Richard H. Swartz, Sandra E. Black, on behalf of the ONDRI Investigators

**Affiliations:** LC Campbell Cognitive Neurology Research, Hurvitz Brain Sciences Program, Sunnybrook Research Institute, University of Toronto, Toronto, Ontario, Canada; Rotman Research Institute, Baycrest, Toronto, ON, Canada; Department of Medical Biophysics, University of Toronto, Toronto, ON, Canada; Centre for Neuroscience Studies, Queens University, Kingston, ON, Canada; Tanz Centre for Research in Neurodegenerative Diseases, University of Toronto, Toronto, ON, Canada; Division of Neurology, Toronto Western Hospital, University Health Network, Toronto, ON, Canada; Stroke Outcomes and Decision Neuroscience Research Unit, Division of Neurology, St Michael’s Hospital, University of Toronto, Toronto, ON, Canada; Thunder Bay Regional Health Research Institute, Thunder Bay, ON, Canada; Department of Medicine (Neurology), Ottawa Hospital Research Instititue and University of Ottawa, Ottawa, ON, Canada; Department of Medical Imaging, University of Toronto, Sunnybrook Health Sciences Centre, Toronto, ON, Canada; Centre for Functional and Metabolic Mapping, Robarts Research Institute, Department of Medical Biophysics, University of Western Ontario, ON, Canada; Department of Medicine (Neurology), Sunnybrook Health Sciences Centre and University of Toronto, ON, Canada

**Keywords:** cortical thickness, quality control, cerebrovascular disease, FreeSurfer, ONDRI, MRI

## Abstract

**Background:** Regional changes to cortical thickness in individuals with neurodegenerative and cerebrovascular diseases can be estimated using specialised neuroimaging software. However, the presence of cerebral small vessel disease, focal atrophy, and cortico-subcortical stroke lesions, pose significant challenges that increase the likelihood of misclassification errors and segmentation failures.

**Purpose:** The main goal of this study was to examine a correction procedure developed for enhancing FreeSurfer’s cortical thickness estimation tool, particularly when applied to the most challenging MRI obtained from participants with chronic stroke and cerebrovascular disease, with varying degrees of neurovascular lesions and brain atrophy.

**Methods:** In 155 cerebrovascular disease patients enrolled in the Ontario Neurodegenerative Disease Research Initiative (ONDRI), FreeSurfer outputs were compared between a fully automated, unmodified procedure and a corrected procedure that accounted for potential sources of error due to atrophy and neurovascular lesions. Quality control (QC) measures were obtained from both procedures. Association between cortical thickness and global cognitive status as assessed by the Montreal Cognitive Assessment (MoCA) score was also investigated from both procedures.

**Results:** Corrected procedures increased ‘Acceptable’ QC ratings from 18% to 76% for the cortical ribbon and from 38% to 92% for tissue segmentation. Corrected procedures reduced ‘Fail’ ratings from 11% to 0% for the cortical ribbon and 62% to 8% for tissue segmentation. FreeSurfer-based segmentation of T1-weighted white matter hypointensities were significantly greater in the corrected procedure (5.8mL vs. 15.9mL, p<0.001). The unmodified procedure yielded no significant associations with global cognitive status, whereas the corrected procedure yielded positive associations between MoCA total score and clusters of cortical thickness in the left superior parietal (p=0.018) and left insula (p=0.04) regions. Further analyses with the corrected cortical thickness results and MoCA subscores showed a positive association between left superior parietal cortical thickness and Attention (p<0.001).

**Conclusions:** These findings suggest that correction procedures that account for brain atrophy and neurovascular lesions can significantly improve FreeSurfer’s segmentation results, reduce failure rates, and potentially increase sensitivity to examine brain-behaviour relationships. Future work will examine relationships between cortical thickness, cerebral small vessel disease, and neurodegenerative disease in the ONDRI study.

## INTRODUCTION

Cortical thickness quantification obtained from magnetic resonance imaging (MRI) can been used to examine regional variations of the cerebral cortex that have been associated with normal ageing and dementia due to neurodegeneration (1–4). Cortical thinning in specific topographical regions of the brain has been used to accurately determine patterns of neurodegeneration in mild cognitive impairment (MCI) (5), Alzheimer’s disease (AD), frontotemporal dementia (FTD) (6–13), Parkinson’s disease (PD) (14–17), amyotrophic lateral sclerosis (ALS) (18–21), and vascular cognitive impairment (22–24).

FreeSurfer (FS) is a neuroimaging software package that includes a widely used surface-based analysis technique that is able to automatically estimate cortical thickness from T1-weighted MRI (25). However, degraded image quality and subtle changes introduced by pathology makes it challenging for FS to achieve accurate and reliable brain extraction and white matter (WM) segmentation (26–31). Although FS provides manual intervention steps to troubleshoot its output (eg. via control points, WM lesion edits, and pial edits), they are labour-intensive. Further, they may introduce user-bias, especially in MRI from individuals with significant brain atrophy, cortical stroke lesions, and cerebral small vessel disease. Previous studies examining FS manual correction approaches found that while manual editing may result in differences in morphometrical estimation between the methods in some brain regions (32–37), sensitivity results are inconsistent at individual or clinical group levels (32–34).

Estimation of cortical thickness in patients with cerebrovascular disease (CVD) can be the most challenging due to corticosubcortical chronic stroke lesions, significant volumes of white matter hyperintensities (WMH), lacunar infarcts, MRI-visible perivascular spaces (PVS), cortical microinfarcts, and the presence of focal brain atrophy. Given that the performance of FS’s tissue classification is highly dependent on a uniform intensity of voxels in a particular brain region and the integrity of the neighbouring voxels, vascular lesions and focal brain atrophy often result in erroneous tissue segmentations, particularly in regions with high surface area and curvature (38). These challenges reduce the accuracy of tissue segmentation, which in turn reduces the accuracy of cortical thickness estimation. Since many age-related neurodegenerative diseases have focal and diffuse brain atrophy that is further exacerbated by comorbid cerebrovascular pathology (39), additional procedures to account for these potentially challenging variations in image contrast are needed.

In this paper, we examined results from a FS correction procedure that was applied to MRI obtained from a heterogeneous CVD cohort with varying degrees cerebral small vessel disease, chronic cortico-subcortical stroke lesions, and brain atrophy.

## MATERIALS AND METHODS

### Study Participants

Participants (N=155) recruited to the CVD cohort of the Ontario Neurodegenerative Disease Research Initiative (ONDRI) (40) (http://ondri.ca) were selected for methodological validation of the FS correction procedure for cortical thickness estimation. The ONDRI study is a multi-modal, multi-site observational research study investigating individuals with neurodegenerative diseases. Study participants were recruited at various health centres across Ontario, Canada: London Health Science Centre and Parkwood Institute in London; Hamilton General Hospital and McMaster Medical Centre in Hamilton; The Ottawa Civic Hospital in Ottawa; Thunder Bay Regional Health Sciences Centre in Thunder Bay; St. Michael’s Hospital, Sunnybrook Health Sciences Centre, Baycrest Health Sciences, Centre for Addiction and Mental Health, and Toronto Western Hospital (University Health Network) in Toronto.

Detailed inclusion and exclusion criteria for the ONDRI CVD participants are previously reported (40,41). Briefly, participants who had experienced a mild to moderate ischemic stroke event, documented with MRI or CT, over 3 months prior to enrollment, a Modified Rankin Scale (MRS) score (42) ranging from 0 to 3, and a Montreal Cognitive Assessment (MoCA) score (43) ranging 18-30 were included. Participants were excluded if they had severe cognitive impairment, aphasia, a non-vascular cause of symptoms, inability to write or had severe functional disability preventing them to perform assessments, a history of dementia prior to the stroke event, had severe claustrophobia or other contra-indications to MRI procedures. Ethics approval was obtained from all participating institutions and performed in accordance with the Declaration of Helsinki. All participants provided informed consent, and subsequently underwent clinical evaluation, MRI, and other assessments as part of the full ONDRI protocol (40).

### MRI Acquisition & Pre-processing

MRI protocols were harmonised with the Canadian Dementia Imaging Protocol (CDIP) (44), and were in compliance with the National Institute of Neurological Disorders and Stroke–Canadian Stroke Network Vascular Cognitive Impairment Harmonization Standards (45). Detailed MRI protocols are reported elsewhere (46,47). In brief, the structural MRI used in the current study include: a high-resolution 3D T1-weighted (T1), an interleaved proton density (PD) and T2-weighted (T2), and a T2 fluid-attenuated inversion recovery (FLAIR) images.

ONDRI’s structural image processing pipeline (47) will be considered as the pre-processing step for the Corrected FS procedure. Briefly, ONDRI’s neuroimaging platform used previously published and validated methods, where outputs were further subjected to comprehensive quality control measures from ONDRI’s neuroinformatic platform using a novel outlier detection algorithm for the identification of anomalous data (48,49). This comprehensive multi-feature segmentation pipeline was applied to co-registered T1, PD, T2, and FLAIR images to generate skull stripped and tissue segmentation masks for each individual, which included manual tracing of cortico-subcortical stroke lesions that were identified and verified on T1 and FLAIR images by an expert research neuroradiologist. The final output of the pipeline produced a skull-stripped brain mask with segmented voxels comprising of 4 different ‘normal tissue’ classes and 5 different ‘lesion tissue’ classes: normal appearing white matter (NAWM), normal appearing gray matter (NAGM), sulcal and ventricular cerebrospinal fluid (sCSF/vCSF), periventricular and deep white matter hyperintensities (pWMH/ dWMH), lacunes, MRI-visible perivascular spaces (PVS), and cortico-subcortical stroke lesions. The skull stripped and lesion-labelled masks were introduced at different processing stages of the Corrected FS procedure described below.

### FreeSurfer (FS) Processing Overview

All scans were processed using the stable version of FS (Linux FSv6.0). Two methods were applied to the same participant’s MRI: a) Unmodified FS and b) Corrected FS. After applying the two methods, visual inspection was performed by two experienced neuroimaging analysts (M.H. = rater1; K.W. = rater2). The images were either rated a “pass” or “fail” based on the overall cortical ribbon and tissue segmentation as described in the Quality Control Assessment Procedures in the following section.

#### Unmodified FS

The unmodified procedure involved the standard reconstruction steps in the FS pipeline with the default settings on all participants without any manual interventions. Briefly, the standard reconstruction steps included skull stripping, WM segmentation, intensity normalisation, surface reconstruction, subcortical segmentation, cortical parcellation and thickness (25).

#### Corrected FS

The corrected procedure involved dividing the reconstruction steps into the following three stages in order to incorporate the skull stripped brain and lesion masks from the ONDRI processing pipeline into FS’s pipeline.

Stage 1 (autorecon1) - This involved replacing the “skull stripped mask” (brainmask.mgz) generated by FS’s standard skull stripping method with an improved skull stripped mask from the ONDRI skull stripping method.

Stage 2 (autorecon2) - The second intervention (autorecon2) involved the integration of lesion masks from ONDRI into the initial version of brain tissue segmentation file (aseg.presurf.mgz) generated by FS’s standard segmentation of the brain which includes subcortical structures, WM, GM, CSF, and white matter hypointensities. The lesions were given an index value of “77” corresponding to the lesion value in the FS pipeline.

Stage 3 - Lastly, the modified aseg.presurf.mgz and brain mask were used as inputs in the last stage of FS pipeline stage 3 (autorecon2-noaseg -autorecon3) for automatic cortical parcellation and statistics.

### Quality Control Assessment Procedures

The accuracy of the cortical ribbon and tissue segmentation from the Unmodified and Corrected FS procedures was evaluated using Freeview, a visualisation tool that is packaged with FS. Using the T1 image as the reference, the cortical ribbon accuracy was assessed visually and given a rating for: 1) Overestimation – areas of non-brain matter, such as the dura mater or skull, were erroneously classified as cortex (**Supplemental Figure S1**); 2) Underestimation – areas of the brain are missing or have been erroneously removed from the brain mask (**Supplemental Figure S2**); 3) Acceptable/Good – no significant areas of over/underestimation; 4) Fail – significant areas of overestimation and/or underestimation, including the complete absence of a cortical ribbon (**Supplemental Figure S3**). Using the T1 GM-WM intensity differences as a reference, the quality of the tissue segmentation (aseg.mgz) was given a PASS/FAIL rating based on the accuracy of WM-GM boundary (FS white matter intensity ~110) (**Supplemental Figure S4**). Two expert neuroimaging analysts performed the visual ratings and achieved high inter-rater reliability results (cortical ribbon: k = 0.9, 95% C.I.: 0.7, 1.00, p < 0.001; tissue segmentation: k = 0.7, 95% C.I.: 0.5, 0.9, p < 0.001).

### Statistical Analysis

Statistical analyses were conducted using Statistical Package for Social Sciences (SPSS v.24) and FS’s packaged analytic software when described. Paired sample t-tests were conducted to determine if the mean lesion volume was significantly different between the unmodified and corrected procedures. This was achieved using the “White matter Hypointensities” identified by FS, which was adjusted for head size using estimated total intracranial volume (eTIV) and log transformed.

A whole brain vertex-wise surface-based cortical thickness analysis was performed on both methods using the built-in general linear model (GLM). Thickness was calculated by the software as the distance between the GM and WM boundaries (also known as the pial surface boundaries) at every vertex in each hemisphere. Each participant’s cortex was anatomically parcellated with every sulcus and gyrus labelled, and resampled to FS’s default average surface map (fsaverage). A 10-mm full-width half-maximum (FWHM) Gaussian spatial smoothing kernel was applied to the surface to improve the signal-to-noise ratio. Age, stroke, and lacunar volumes were included as nuisance regressors. Stroke and lacunar volumes were head size corrected using total intracranial volume.

MoCA total score was included as a regressor of interest to determine the association between cortical thickness and global cognitive status in participants with CVD. Associations between cortical thickness and cognition were further explored using MoCA sub-scores (Visuospatial / Executive, Naming, Memory, Attention, Language, Abstraction, Delayed Recall, and Orientation). Monte Carlo simulations with 5000 iterations were used to correct for multiple comparisons. This method implemented a cluster-wise threshold of 2 and cluster-wise probability (p(cwp)) of p < 0.05 (two-sided). Bonferroni correction was applied across the two hemispheres.

## RESULTS

Study participant demographics and clinical characteristics are summarized in **Table 1**. Quality control (QC) results are summarized in **Table 2**.

**Table 1.** Study participant demographics and neuroimaging volumetrics (n=155). All data are presented as mean (SD) unless otherwise indicated.

| **Demographics** |  |
| --- | --- |
| Age (years) | 69.35 (7.36) |
| Sex, n (%) female | 48 (31) |
| Education, years | 14.69 (2.88) |
| Modified Rankin Scale | 1.01 (0.83) |
| Montreal Cognitive Assessment | 25.29 (2.99) |
| **Neuroimaging Volumetrics, mm<sup>3</sup>** |  |
| White matter hyperintensities | 10167.5 (12837.2) |
| Lacunes | 385.1 (766.7) |
| Enlarged perivascular spaces | 80.0 (139.7) |
| Cortico-subcortical Stroke Lesions | 6785.0 (17317.8) |

**Table 2.** Quality control results for FreeSurfer Unmodified and Corrected procedures.

| Description | Unmodified | Corrected |
| --- | --- | --- |
| **Cortical Ribbon** |  |  |
| Over-estimation | 26% | 0% |
| Under-estimation | 45% | 24% |
| Acceptable | 18% | 76% |
| Fail | 11% | 0% |
| **Tissue Segmentation** |  |  |
| Pass | 38% | 92% |
| Fail | 62% | 8% |

For the cortical ribbon QC, compared to the Unmodified FS procedure, the Corrected ‘Acceptable’ ratings increased from 18% to 76%. For tissue segmentation QC, compared to the Unmodified FS procedure, the Corrected procedure’s ‘Acceptable’ ratings increased from 38% to 92%. For the cortical ribbon QC, the ‘Fail’ ratings were reduced from 11% (Unmodified) to 0% (Corrected). While for the tissue segmentation QC, the ‘Fail’ ratings were reduced from 62% to 8% for Unmodified and Corrected procedures respectively (e.g. **Figure 1**).

**Figure 1.**
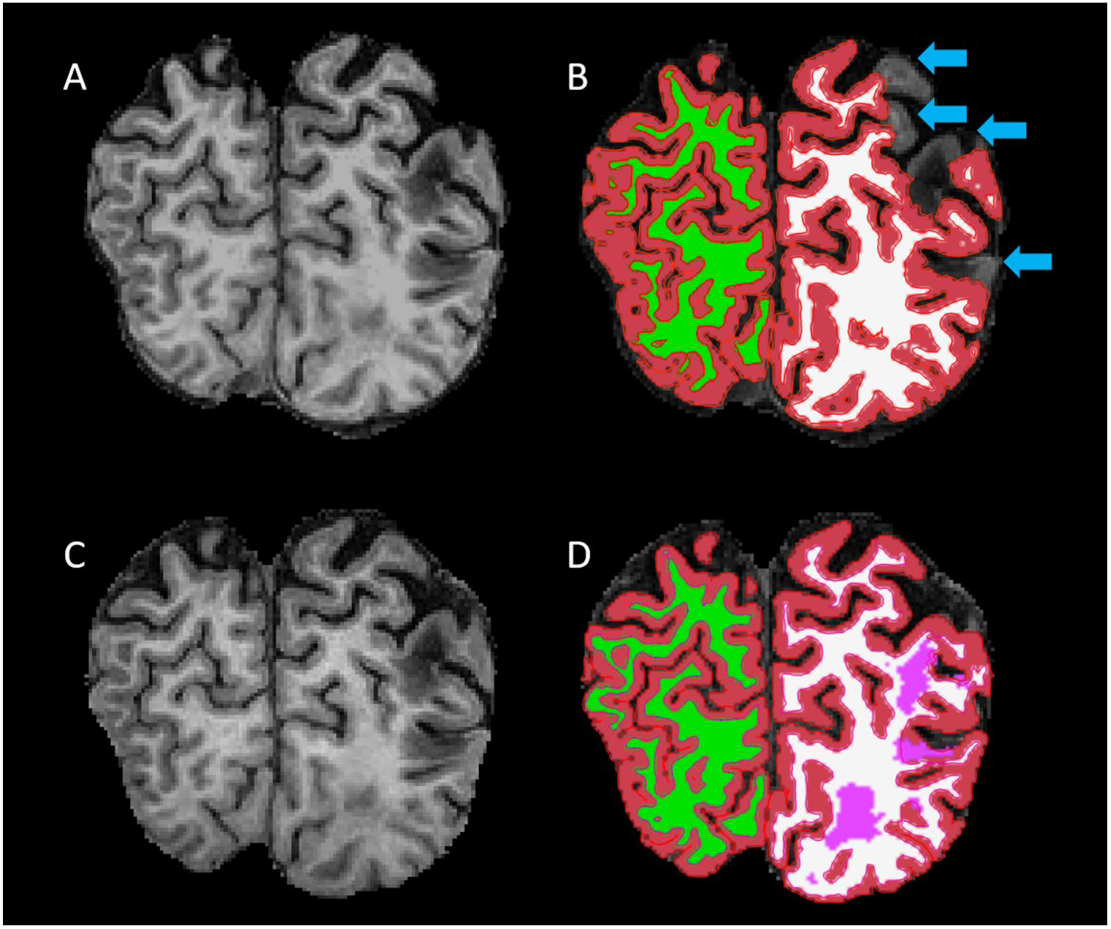
Comparison of outputs generated from the unmodified (A-B) vs the corrected (C-D) FreeSurfer procedures. A) Skull-stripped coronal image from the unmodified procedure. B) Segmentation result from the unmodified procedure overlaid on the skull-stripped T1. C) Skull-stripped coronal image from the corrected procedure. D) Segmentation result from the unmodified procedure overlaid on the skull-stripped T1. Segmentation: Red = gray matter; Green/White = right/left white matter; Pink = lesion. Blue arrows point to areas of the brain that are missing or have been erroneously removed from the segmentation.

When comparing Unmodified and Corrected procedures, results from a paired sample t-test revealed a significant increase (~63%) in eTIV-adjusted log white matter hypointensity volumes, (5824.5 ± 6378.4 mm^3^ to 15877.1 ± 17964.2 mm^3^, p < 0.001).

Cortical thickness analyses based on Unmodified FS revealed no significant associations with MoCA total score after accounting for age, stroke, and lacunar volumes. However, the same analyses based on the Corrected FS revealed significant clusters in the left superior parietal and left insula regions were positively associated with MoCA total score (p_(cwp)_ = 0.018; p_(cwp)_ = 0.040, respectively) (**Table 3**, **Figure 2**). Further analyses with the Corrected data and MoCA sub-scores using the significant clusters showed a positive association between left superior parietal thickness and the Attention sub-score.

**Table 3.**
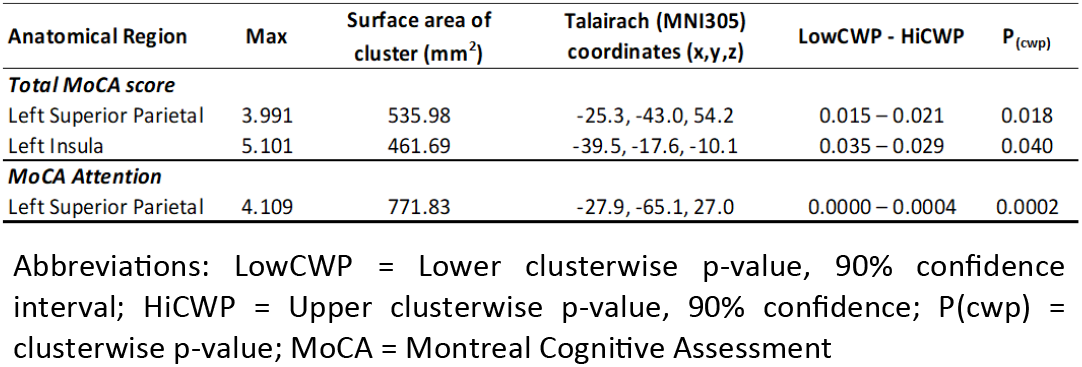
Cortical thickness analyses showing significant clusters with Montreal Cognitive Assessment

**Figure 2.**
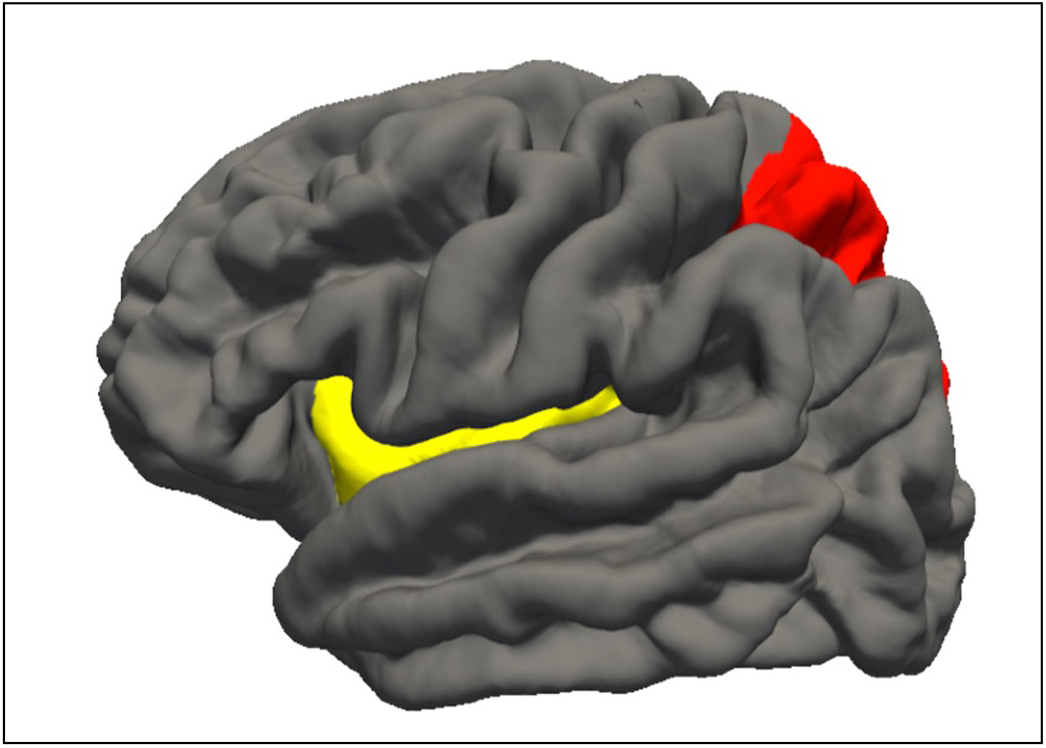
Cortical thickness regions showing significant associations with cognition. Red = left superior parietal; Yellow = left insula. Both regions were associated with general cognition and the left superior parietal was associated with attention.

## DISCUSSION

Regional cortical thickness measures obtained from participants’ MRI using FS is a useful imaging biomarker of cortical atrophy, within and between the various disease cohorts represented in ONDRI. The increase in accuracy and reduction in failure rates due to our correction procedures described here has the opportunity to advance the study of structural biomarkers in neurodegeneration, by minimising data loss and increasing statistical power. This correction procedure enabled the investigation of participants with significant atrophy and cerebrovascular lesion burden, which can present significant challenges to cortical thickness estimation, cortical and subcortical volumetrics, and other downstream processes (e.g. connectivity analyses of functional and diffusion MRI). Moreover, the correction procedures may improve the sensitivity of estimated features that may have otherwise been undetectable.

In the unmodified procedure, a failure rate of more than 60% was reported for tissue segmentation. This is in line with the concept that most segmentation difficulties reported in individuals with CVD result from inaccurate identification of tissue boundaries, which is highly dependent on the homogenous intensity of voxels in a particular brain region, especially in those with high surface area and curvature (50). Accurate and reliable skull stripping is important for cortical thickness estimation, since false positive classification of non-brain tissue (e.g. skull, dura and pial maters) could result in poor estimation of the GM-WM border, which in turn can result in erroneous patterns of cortical thickness. Skull stripping segmentation accuracy is particularly relevant in ageing and neurodegenerative populations, where brain atrophy is accompanied by increased CSF volumes and a decreased separation between GM and WM intensities (10,14,15,18,22,51).

While small acute strokes may have minimal effects on tissue segmentation, large chronic cortico-subcortical stroke lesions introduce alterations to brain morphometry resulting in failed segmentation in most brain segmentation algorithms (50,52–54). Although this issue is particularly relevant in individuals with CVD, cerebral small vessel disease and brain atrophy that are commonly observed in patients with Alzheimer’s and other related dementias present similar challenges when estimating cortical thickness.

Incorporating more accurate brain extraction and lesion masks from ONDRI reduced the overall failure rate to less than 8% when the corrected procedure was applied. This improvement could be attributed to the use of multi-modal imaging sequences in the ONDRI structural neuroimaging pipeline (47). This method produces consistent and accurate brain extraction and lesion segmentation. Although imaging markers of small vessel disease, such as WMH, appear hyperintense (bright) on PD/T2 and FLAIR MRI, these lesions appear hypointense or isointense to GM on T1, thus overlapping in intensity with normal appearing GM (55). If present in confluent patches, it can result in significant inflation of GM voxel misclassification when using only T1-based segmentation approaches (56). Considering the significant WMH burden and atrophy in our sample, it was helpful that the FS pipeline allowed for these types of interventions. In line with this, we found a significant increase in white matter hypointensities burden (~63%) after incorporating ONDRI’s lesion segmentation to the FS pipeline.

Several studies have underscored the importance of optimal lesion segmentation in various clinical population (57–61), particularly in populations at risk of developing small vessel disease (59,62–64). A recent systematic review by Frey et al., (2019) provided a comprehensive overview of the importance of WMH segmentation in large-scale MRI studies. They proposed a clear need for developing a guideline to cover the description of WMH segmentation approach, as a way of optimising the multitude of segmentation techniques available. This is crucial, especially in medium to large sample size studies with clinical populations that donate their time to research. Furthermore, the flexibility of the FS pipeline to allow for such modification supports the individualised imaging methods used in the ONDRI study. This increases the study’s statistical power whilst including participants with challenging pathologies that otherwise might have failed when processed using the default settings, and in turn, reduces sampling bias related to the imaging method requirements (27).

Only data that underwent the FS correction demonstrated a relationship with cognition, whereby greater corrected cortical thickness in the left superior parietal cortex and in the insula was associated with higher MoCA total scores. Further analysis with MoCA sub-scores revealed that corrected cortical thickness in the left superior parietal cortex was associated in particular with higher Attention sub-scores. Several studies have reported a significant association between cortical thickness and cognitive function in participants with SVD and other diseases associated with vascular risk factors (66–70). Across these studies the effect of cortical thickness varies, with some reporting relationships with executive function, processing speed, memory (66,71), whilst others reporting relationships with memory and attention (67,70). A study by Hilal et al. (2015) demonstrated that WMH and microbleeds were associated with thinning in the temporal and insular regions and associated multi-domain cognitive dysfunction. The insula is an important structure with extensive connections to cortical and subcortical regions, and is involved in various processes, such as empathy, emotion, body sensation, and other aspects of social cognition (73,74). Thus, insular atrophy as a result of stroke could lead to significant cognitive dysfunction and socioemotional deficits in participants with cerebral small vessel disease and other comorbid neurodegenerative diseases (75–77). Further, the observed association between superior parietal thickness and the Attention sub-score is consistent with recent work showing that smoking-related superior parietal thinning was associated with decreased global cognition, as well as decreased visuospatial and attentional functioning (78). This is in line with the concept that better vascular health is associated with increased superior parietal thickness in neurodegenerative diseases (79–81), suggesting a compensatory response to early brain pathological changes (82). Future analyses using our method will investigate the associations between vascular risk factors and cortical thickness in predicting cognitive decline in neurodegenerative diseases with comorbid cerebral small vessel disease.

The ability to decrease the failure rate was the key strength of this work. Although our correction procedures were derived from ONDRI’s imaging pipeline, similar correction procedures that can account for vascular lesions and brain atrophy could be applied in other studies using FS (or any number of cortical thickness estimation tools) to study challenging clinical populations (57,83–85). Hence, the decision to validate and apply this method to individuals with CVD presenting with a range of various combined pathologies including focal and global atrophy, large and small cortico-subcortical chronic stroke lesions, diffuse and focal WMH, lacunar infarcts, cortical microinfarcts, and enlarged PVS (55,86). This combination of brain pathologies brings a unique set of potential challenges for cortical thickness estimation.

The findings reported here should also be considered in light of several limitations. The cross-sectional analysis of this project limits our ability to examine the robustness of our method longitudinally. As ONDRI is a longitudinal study, future work will implement our method at several follow-up time points, within and between all disease cohorts, providing a unique opportunity to investigate relationships between cortical thickness and other neurodegenerative biomarkers for predicting disease progression. Another benefit to the FS correction is its potential to facilitate better understanding of brain-behaviour relationships by increasing the sensitivity and accuracy of the cortical estimation tool. As demonstrated, only corrected cortical estimations correlated with a measure of global cognitive status. Future work will examine cross-sectional and longitudinal relationships between cortical thickness, vascular risk factors, neurodegeneration, and associations with comprehensive neuropsychological testing (87).

## CONCLUSIONS

Given these results, our findings strongly suggest that individualised accounting of brain atrophy and neurovascular lesions in cortical thickness estimation tools such as FS, can significantly improve the segmentation results, reduce failure rates to minimise biased samples, and potentially increase sensitivity to examine brain-behaviour relationships. Most importantly, these correction efforts invested to reduce data loss and inaccuracies, acknowledge the significant time and effort our patients have donated to participate in the ONDRI research study.

## Acknowledgements

We would like to thank the ONDRI participants for the time, consent, and participation in our study. Thank you to the L.C. Campbell Foundation, and the analysts and software developers in the LC Campbell Cognitive Neurology research team who have contributed to the ONDRI imaging analysis, including Edward Ntiri, Parisa Mojiri, Rita Meena, and Pugaliya Puveendrakumaran.

This paper is available in preprint version online: https://doi.org/10.1101/2020.08.04.236760

## Author Contributions

MO: Conceptualisation, Data Curation, Formal Analysis, Investigation, Methodology, Project Administration, Software, Validation, Visualisation, and Writing (draft, review, and editing)

JR: Conceptualisation, Data Curation, Formal Analysis, Investigation, Methodology, Software, Validation, Visualisation, Writing (draft, review, and editing), and Supervision

PRR: Data Curation, Formal Analysis, and Writing (review and editing)

MFH: Data Curation, Validation, Visualisation, and Writing (review and editing)

KW: Data Curation, Validation, Visualisation, and Writing (review and editing)

CJMS: Data Curation, Project Administration, and Writing (review and editing)

MG: Writing (review and editing)

DK: Data Curation, Project Administration, Writing (review and editing)

MCT: Writing (review and editing), Supervision, Funding Acquisition

DB: Data Curation, Software, Writing (review and editing)

GS: Resources, Writing (review and editing)

AH: Resources, Funding Acquisition

JLD: Resources, Data Curation, Writing (review and editing)

DD: Resources, Data Curation, Funding Acquisition

SCS: Data Curation, Resources, Funding Acquisition

SS: Data Curation, Supervision

RB: Data Curation, Resources, Supervision, Funding Acquisition

RHS: Data Curation, Writing (review and editing), Resources, Supervision, Funding Acquisition

SEB: Conceptualisation, Methodology, Supervision, Writing (review and editing), Resources, Funding Acquisition

## ONDRI Investigators

Michael Strong, Peter Kleinstiver, Natalie Rashkovan, Susan Bronskil, Michael Borrie, Elizabeth Finger, Corinne Fischer, Andrew Frank, Morris Freedman, Sanjeev Kumar, Stephen Pasternak, Bruce Pollock, Tarek Rajji, Dallas Seitz, David Tang-Wai, Brenda Varriano, Agessandro Abrahao, Marvin Chum, Christen Shoesmith, John Turnbull, Lorne Zinman, Julia Fraser, Bill McIlroy, Ben Cornish, Karen Van Ooteghem, Frederico Faria, Manuel Montero-Odasso, Yanina Sarquis-Adamson, Alanna Black, Barry Greenberg, Wendy Hatch, Chris Hudson, Elena Leontieva, Ed Margolin, Efrem Mandelcorn, Faryan Tayyari, Sherif Defrawy, Don Brien, Ying Chen, Brian Coe, Doug Munoz, Alisia Bonnick, Leanne Casaubon, Dar Dowlatshahi, Ayman Hassan, Jennifer Mandzia, Demetrios Sahlas, Gustavo Saposnik, David Breen, David Grimes, Mandar Jog, Anthony Lang, Connie Marras, Mario Masellis, Tom Steeves, Dennis Bulman, Allison Ann Dilliott, Mahdi Ghani, Rob Hegele, John Robinson, Ekaterina Rogaeva, Sali Farhan, Hassan Haddad, Nuwan Nanayakkara, Courtney Berezuk, Sabrina Adamo, Mojdeh Zamyadi, Stephen Arnott, Brian Tan, Malcolm Binns, Wendy Lou, Kelly Sunderland, Athena Theyers, Abiramy Uthirakumaran, Guangyong (GY) Zou, Sujeevini Sujanthan, Mojdeh Zamyadi, David Munoz, Roger A. Dixon, John Woulfe, Brian Levine, Paula McLaughlin, JB Orange, Alicia Peltsch, Angela Roberts, Angela Troyer.

## SUPPLEMENTAL FIGURES

**Figure S1.**
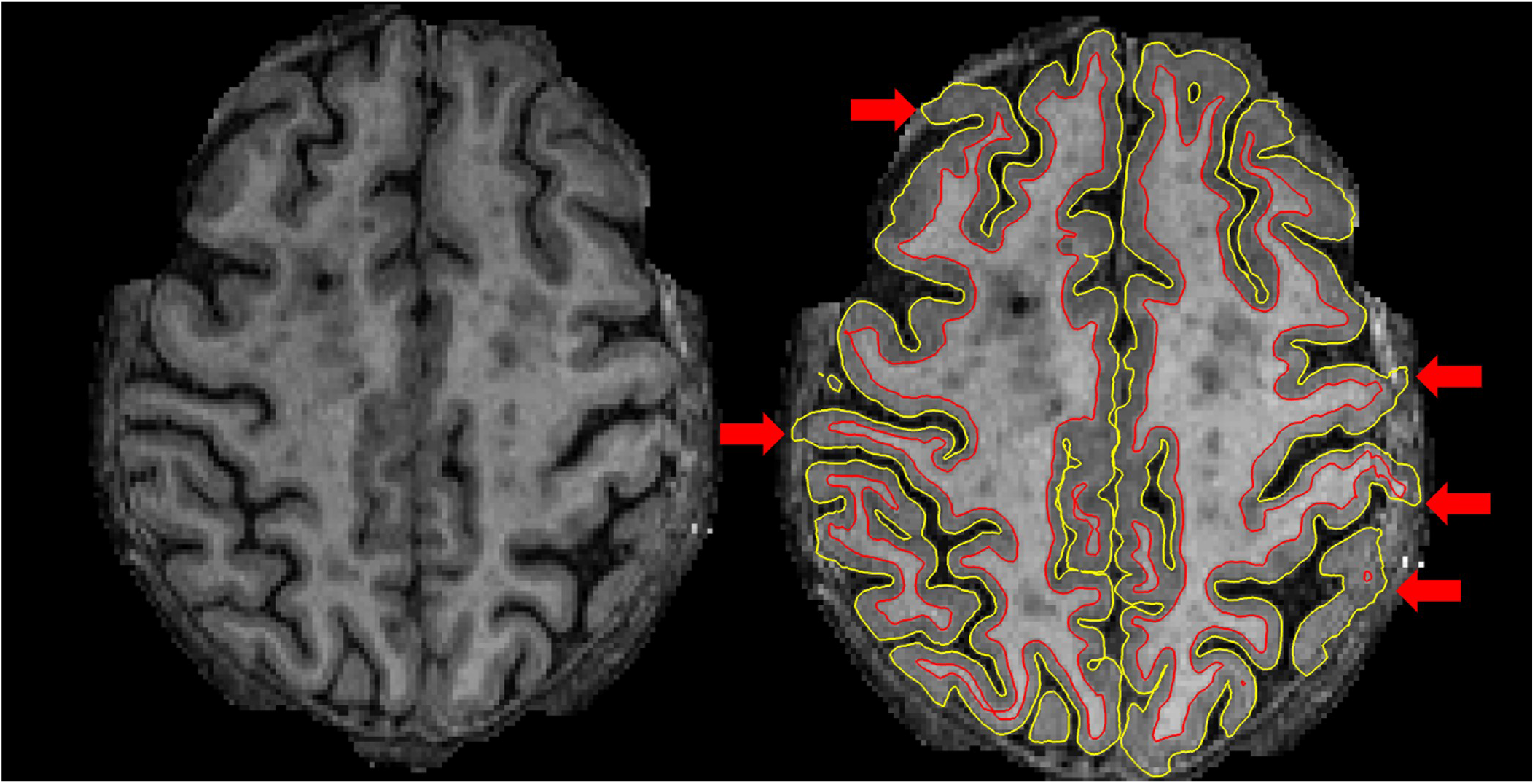
Cortical ribbon Quality Control ‘Overestimation’ rating example. Left image shows an axial slice of a T1 that was skull-stripped using the Unmodified FreeSurfer procedure. Right image shows the segmentation that results from the Unmodified procedure. Red arrows point to areas of non-brain matter, such as the dura or skull, that were erroneously classified as cortex and/or white matter.

**Figure S2.**
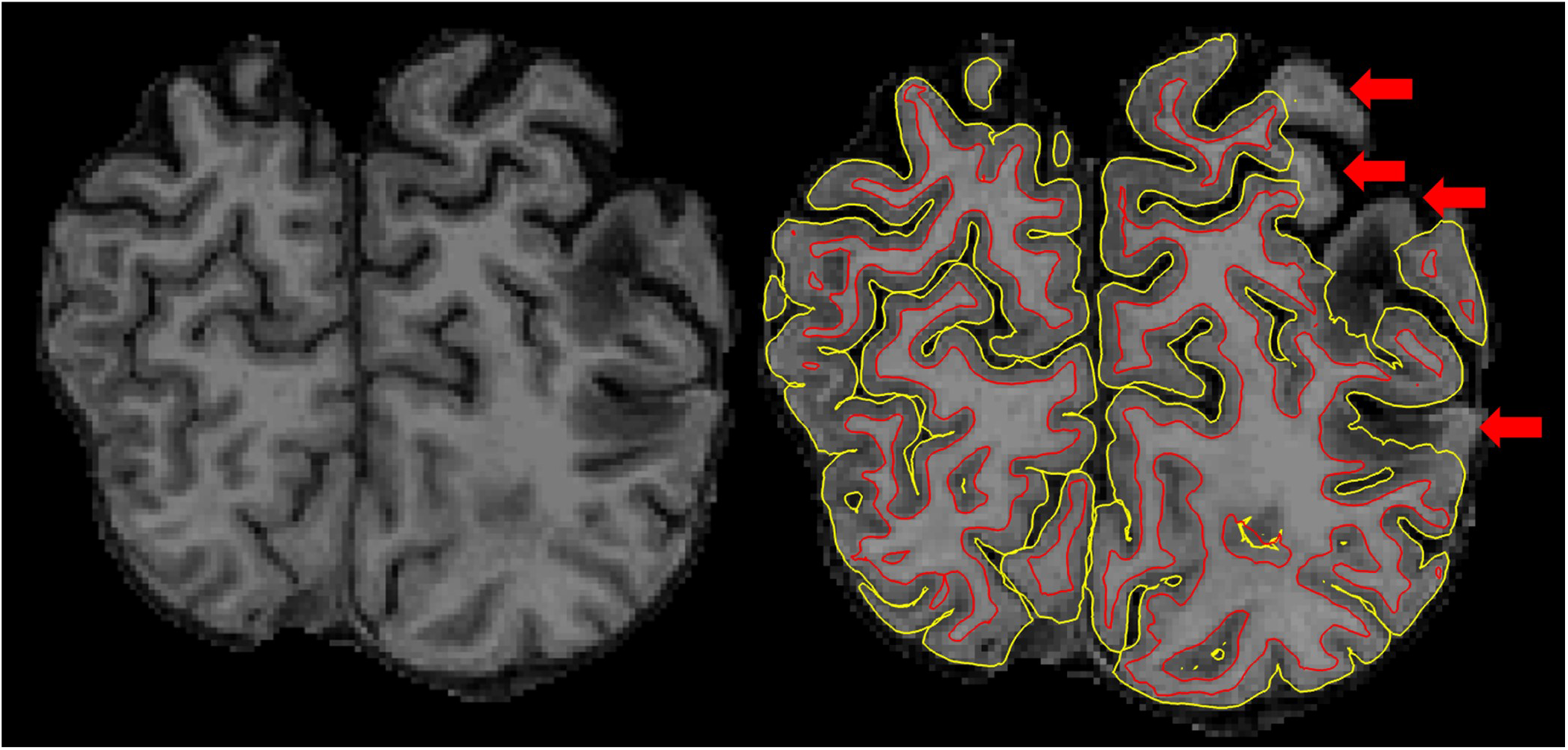
Cortical ribbon Quality Control ‘Underestimation’ rating example. Left image shows a coronal slice of a T1 that was skull-stripped using the Unmodified FreeSurfer procedure. Right image shows the segmentation (ribbon) that results from the Unmodified procedure. Red arrows point to areas of the brain that are missing or have been erroneously removed.

**Figure S3.**
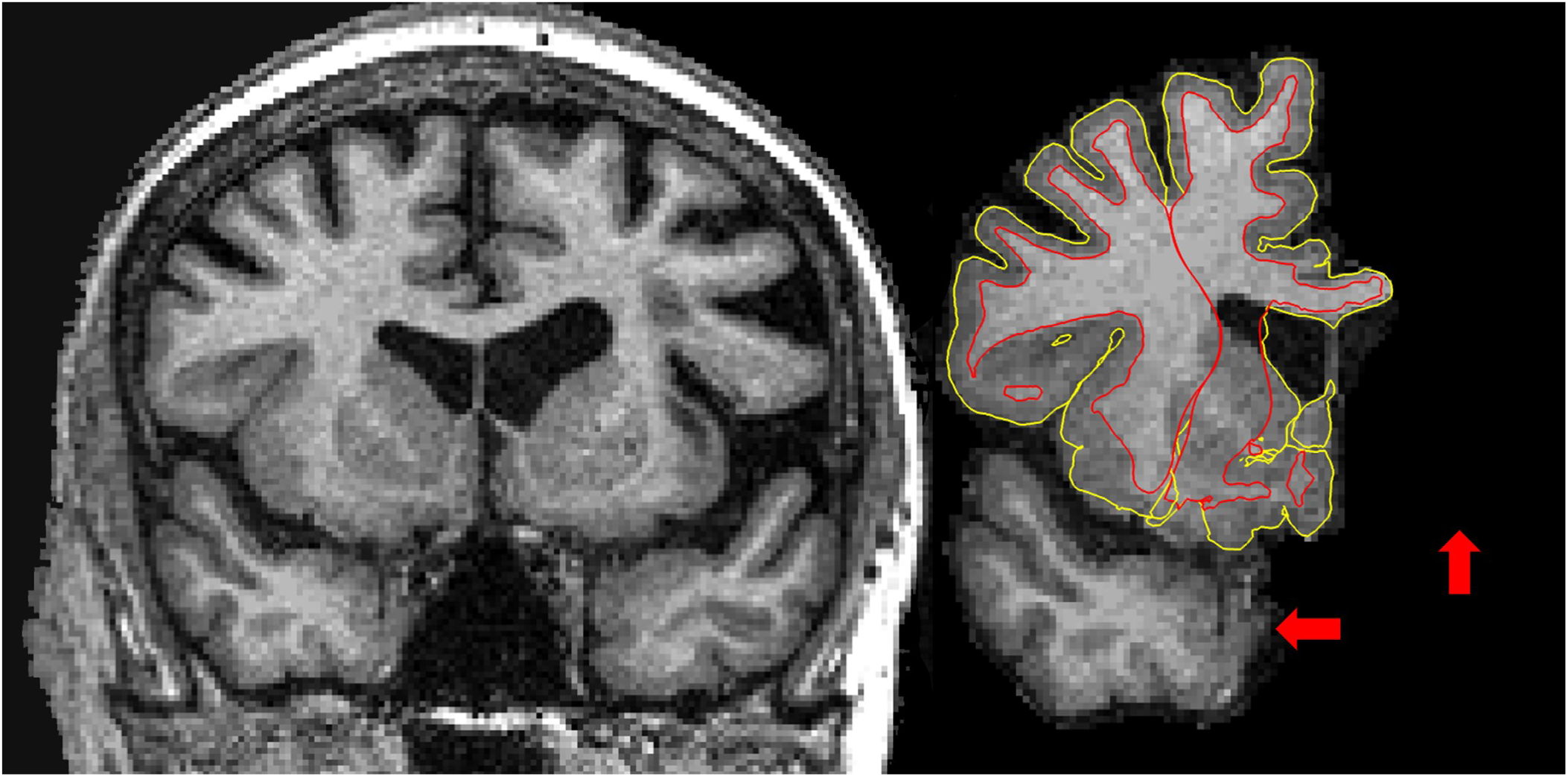
Cortical ribbon Quality Control ‘Fail’ rating example. Left image shows a coronal slice of the original T1 that was run through the Unmodified FreeSurfer procedure. Right image shows the segmentation (ribbon) that resulted from the Unmodified procedure. Red arrows point to areas of the brain that are missing and have been erroneously removed (temporal lobe and one entire hemisphere).

**Figure S4.**
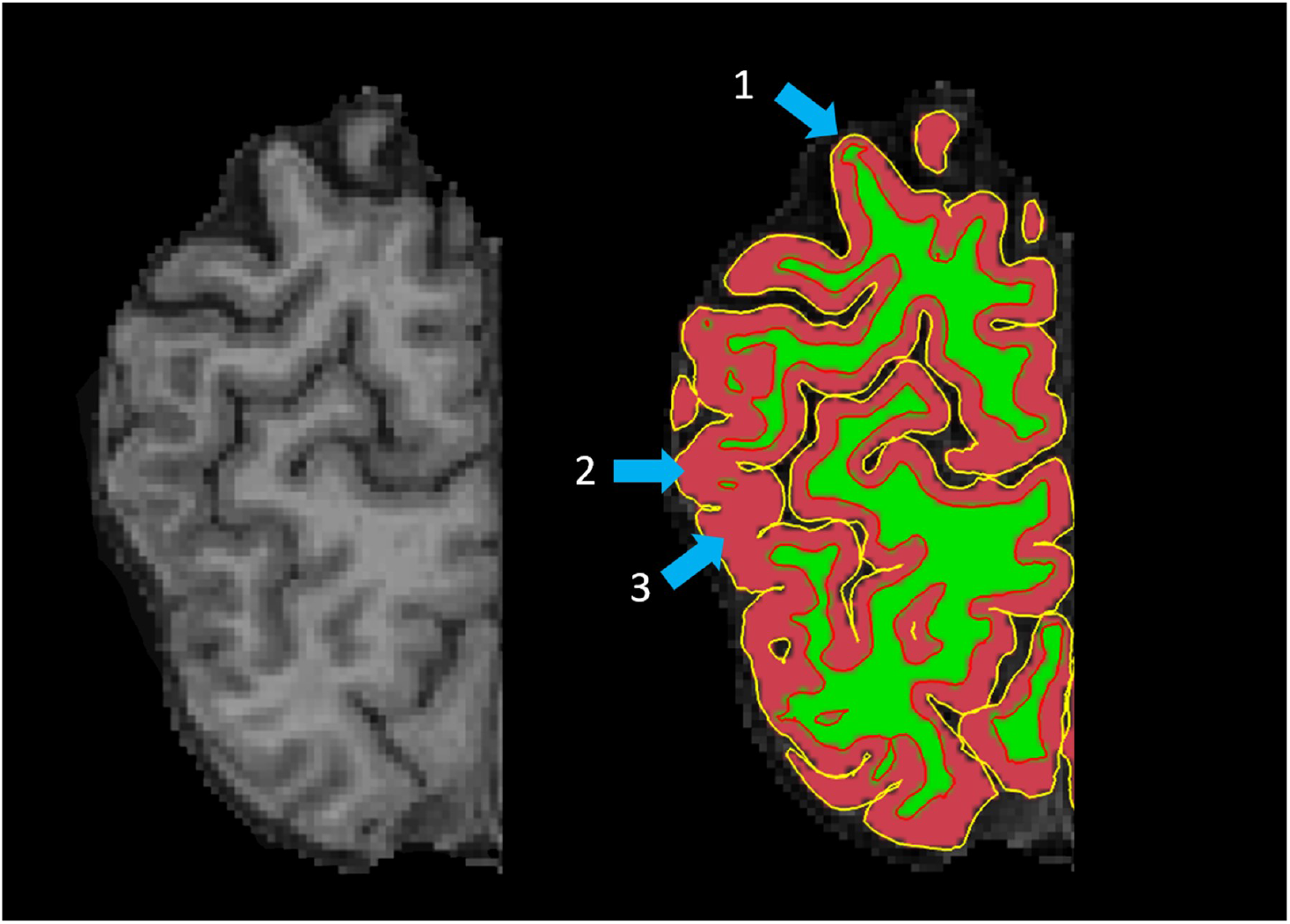
Tissue segmentation Quality Control ‘Fail’ rating example. Left image shows a coronal slice of the original T1 that was run through the Unmodified FreeSurfer procedure. Right image shows the tissue segmentation that resulted from the Unmodified procedure where white matter (WM) is shown in green and gray matter (GM) is shown in red. Blue arrows point to areas where: 1) GM is misclassified as WM, underestimating cortical thickness, 2-3) WM is misclassified as GM, overestimating cortical thickness.

